# Robotic microscopy for everyone: the OpenFlexure Microscope

**DOI:** 10.1101/861856

**Authors:** Joel T. Collins, Joe Knapper, Julian Stirling, Joram Mduda, Catherine Mkindi, Valeriana Mayagaya, Grace A. Mwakajinga, Paul T. Nyakyi, Valerian L. Sanga, Dave Carbery, Leah White, Sara Dale, Zhen Jieh Lim, Jeremy J. Baumberg, Pietro Cicuta, Samuel McDermott, Boyko Vodenicharski, Richard Bowman

## Abstract

Optical microscopes are an essential tool for both the detection of disease in clinics, and for scientific analysis. However, in much of the world access to high-performance microscopy is limited by both the upfront cost and maintenance cost of the equipment. Here we present an open-source, 3D-printed, and fully-automated laboratory microscope, with motorised sample positioning and focus control. The microscope is highly customisable, with a number of options readily available including trans- and epi-illumination, polarisation contrast imaging, and epi-florescence imaging. The OpenFlexure Microscope has been designed to enable low-volume manufacturing and maintenance by local personnel, vastly increasing accessibility. We have produced over 100 microscopes in Tanzania and Kenya for educational, scientific, and clinical applications, demonstrating that local manufacturing can be a viable alternative to international supply chains that can often be costly, slow, and unreliable.

## 1 Introduction

For centuries optical microscopy has been the foundation of scientific imaging and analysis in many aspects of medicine, life and physical sciences [1]. Commercially available high-end microscopes can provide precise motorised high-resolution imaging capabilities, however these are often prohibitively expensive. This is compounded in the developing world where unreliable supply chains inflate prices, and slow acquisition [2]. Maintaining equipment adds further burden as devices have often not been designed for these operating environments [3], and neither manufacturers representatives nor spare parts are available locally, leading to much of the laboratory equipment in resource-poor countries being out-of-service [4].

Open-source hardware is poised to revolutionise the distribution of scientific instrumentation, impacting research, local manufacturing, and education [5, 6]. Open science hardware designs are already in use for both research and education across a wide range of disciplines [7, 8, 9, 10, 11]. Within research laboratories in particular, 3D printers have become an increasingly common item of equipment. This has led a number of open science hardware projects to use 3D printing as an accessible platform for locally prototyping and manufacturing laboratory-grade devices [10, 11, 12].

Here we present the OpenFlexure Microscope (OFM), a 3D-printed, laboratory-grade automated microscope. The OFM aims to make automated microscopy accessible both to projects with a modest budget in established labs, and to developing nations aiming to adopt more advanced scientific research technologies. It is built around a well-characterised flexure mechanism [13], providing precise 3D motion for focus and sample positioning. The range of mechanical motion is smaller than traditional mechanical stages, with 12 × 12 × 4 mm travel, and due to the primarily plastic construction, our design has a limited load capacity unsuitable for very large or heavy samples. However, both the range of motion, and load capacity, are ample for most microscopy applications. It can be constructed with a range of different interchangeable optics modules, enabling different imaging modalities (Section 2.1) and allowing different cameras and imaging lenses to be used for applications from blood smear parasitology to school teaching.

The OFM has now been trialled in a range of applications, and reproduced by a number of groups around the world. The design (Figure 1) has been refined to reduce assembly time, ease component sourcing, and maximise reliability. These ongoing optimisations are key to a sustainable and useful open hardware project that is trusted for critical applications. Version 6.0 of the OpenFlexure Microscope represents a well tested microscopy platform, enabling the rapid prototyping of new instrumentation, and replication of high quality research tools, even in resource-poor settings.

**Figure 1:**
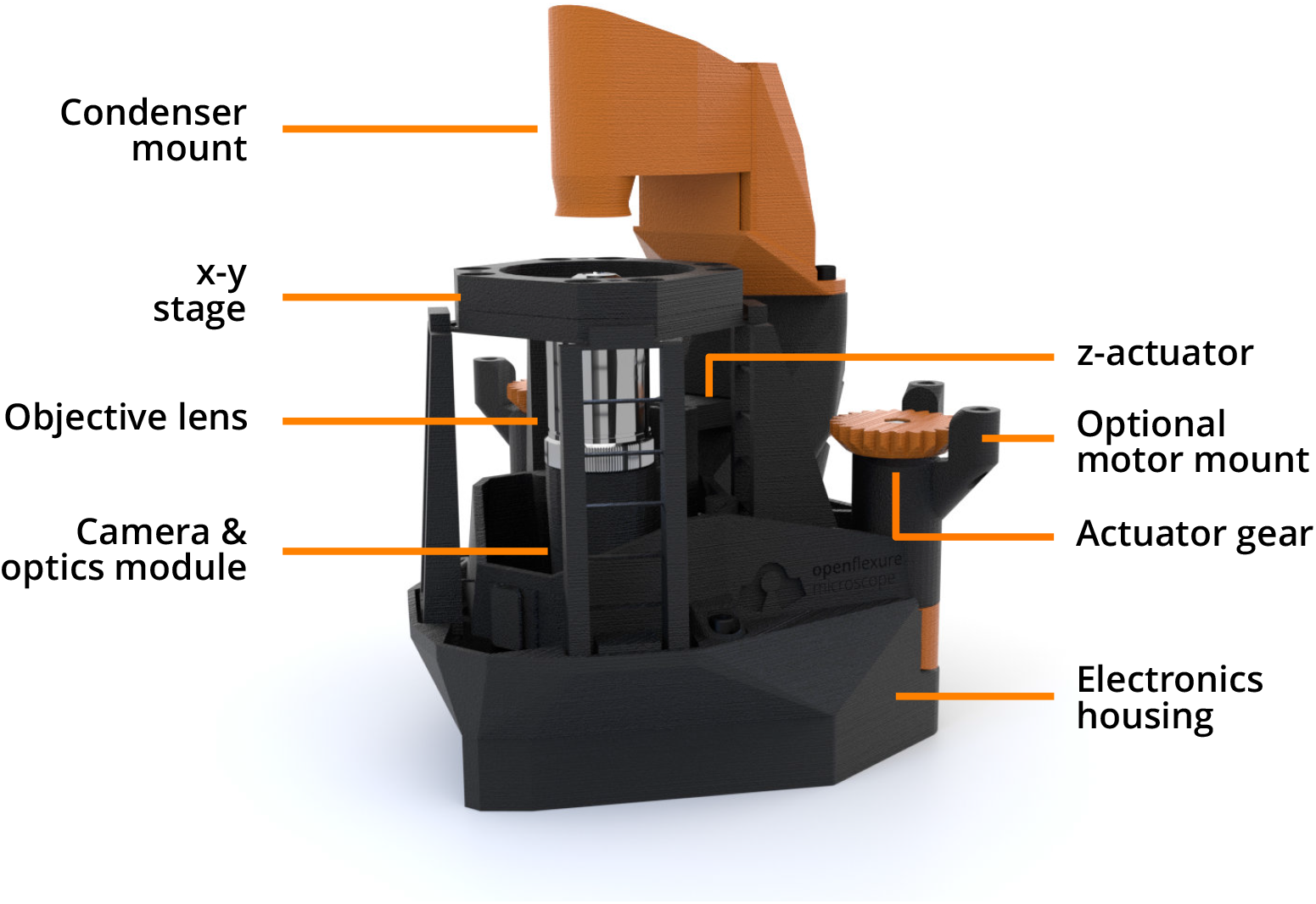
Overview of the OpenFlexure Microscope design, in transmission bright-field configuration. The condenser mount houses an illumination LED and a plastic condenser lens, while the optics module sits below the stage and houses an objective lens, tube lens, and camera. The entire optics module is attached to the *z*-actuator, providing variable focus. Both the optics module and *x-y* stage are controlled by actuator gears at the back of the microscope, optionally driven by stepper motors. A detachable electronics housing stores optional electronic parts, such as motor controllers and a Raspberry Pi, for automation.

## 2 Imaging capabilities

The OpenFlexure Microscope uses an interchangeable optics module to easily change between imaging modes. Most of the optics modules use standard RMS objectives to yield high-quality images expected from a laboratory microscope, and an option is available to use an inverted, spatially offset webcam lens to provide magnification at a low-cost for educational use. Optics modules are available for bright-field (with both trans- and epi-illumination) imaging, polarisation contrast imaging, and epifluorescence imaging. The estimated parts costs for various configurations of the microscope are provided in Supporting Information 1.

### 2.1 Imaging modes

#### 2.1.1 Bright-field trans-illumination

Bright-field trans-illumination is the “standard” imaging mode for the OFM. White light illumination, provided by a diffused LED, is focused onto the sample using a condenser lens. Light passing though the sample is imaged with an RMS objective, a tube lens, and an 8MP CMOS sensor (Raspberry Pi camera V2) (Figure 2a). Both finite and infinite conjugate objectives can be used by varying the position of a tube lens. Figure 2a shows an image of a Giemsa-stained blood smear, obtained at the Ifakara Health Institute, for the purpose of malaria diagnosis. The image is obtained with a 100×, 1.25NA oil immersion objective. Individual red blood cells, with dimensions on the order of microns, are clearly resolved. A ring-form trophozoite of *Plasmodium falciparum* within a red blood cell is highlighted in the figure inset (contrast has been digitally enhanced for clarity by increasing brightness and gamma). Despite being only 1 μm to 2 μm across, the morphology of these ring-form parasites is also clearly resolved by the OFM.

**Figure 2:**
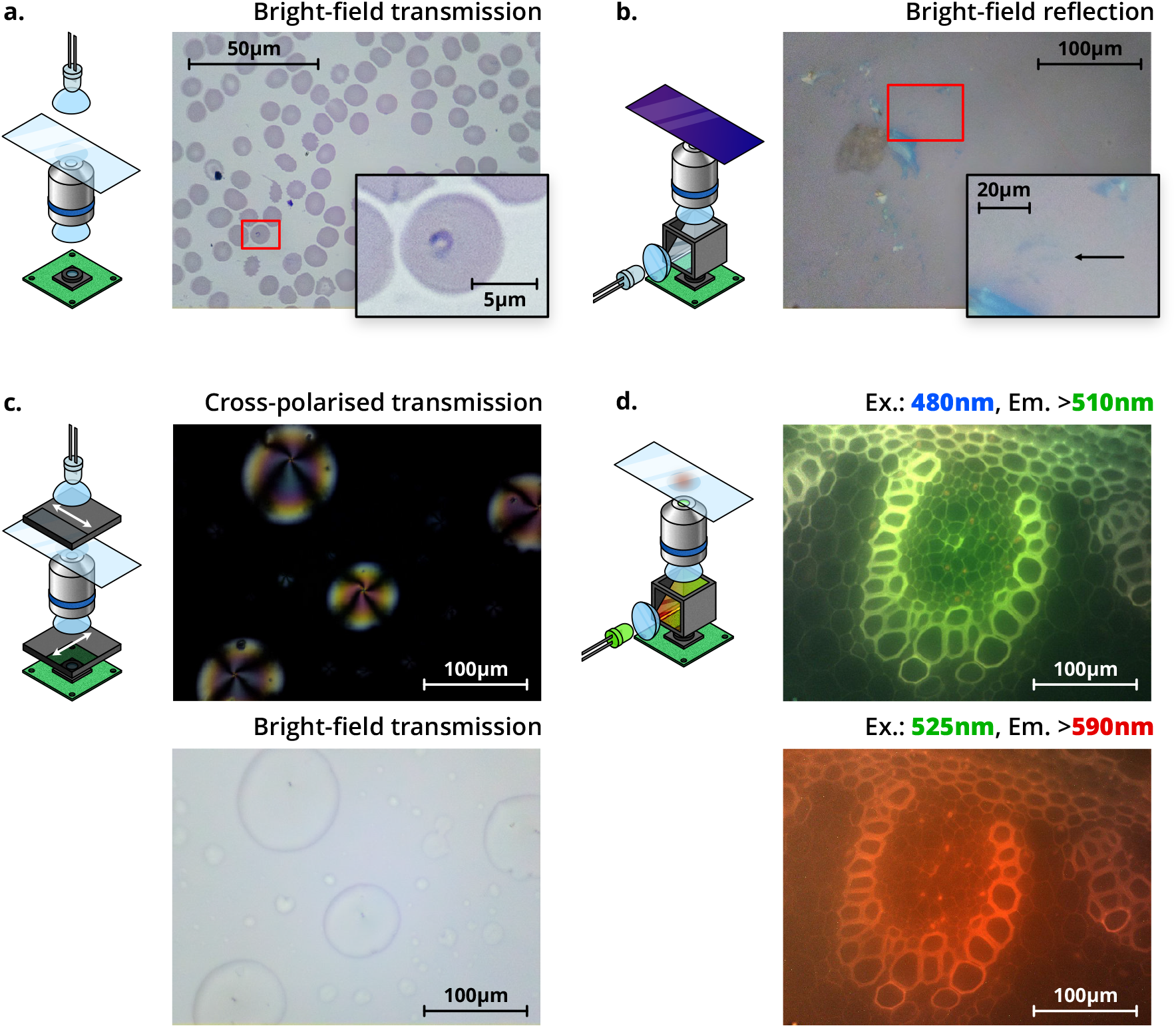
Schematics (left) and images (right) of four different imaging modalities possible using the OFM. **a.** Trans-illumination bright-field imaging of a Giemsa-stained thin blood smear, obtained with a 100×, 1.25NA oil immersion objective. The inset shows a magnified section of the image, highlighting a ring-form trophozoite of *Plasmodium falciparum*. **b.** Epi-illumination bright-field imaging of a group of thin graphene flakes, obtained with a 40×, 0.65NA air objective. The inset shows a magnified section of the image, highlighting a resolvable tri-layer graphene flake (contrast has been digitally enhanced for clarity by increasing brightness and gamma). **c.** Polarisation-contrast trans-illumination image of 5CB liquid crystal droplets. A bright-field image is shown below for comparison. Both images were obtained with a 40×, 0.65NA air objective, and depict the same region of the sample. Arrows on the polarisers denote the transmission axis. **d.** Fluorescence images of unstained Lily of the Valley (*convallaria majalis*) rhizome, at two different wavelengths. The excitation wavelength (“Ex.”), and minimum emission wavelength imaged (“Em.”) are shown above the respective panels. Both images were obtained with a 40×, 0.65NA air objective, and depict the same region of the sample. Both the colour of illumination LED, and the filters in the filter cube, will change depending on the application.

Image quality is in line with expectations for manual microscopes currently used for malaria diagnosis. However, digital images make it possible to re-examine a slide without physically preserving and transporting it. This improves record-keeping, quality control, and training of both new and experienced technicians. Automatic acquisition of images (Section 2.3) goes further, improving throughput of samples and paving the way for automated image processing to assist diagnostic technicians.

#### 2.1.2 Bright-field epi-illumination

Reflection (epi-) bright-field imaging is enabled by inserting a 50/50 beam splitter at 45° within a printed filter cube between the tube lens and the sensor. Illumination is provided by a collimated diffused LED reflected through the objective by the beamsplitter (schematized in Figure 2b). Light reflected from the sample is partially transmitted through the beam splitter and onto the sensor, enabling imaging of opaque samples.

Figure 2b shows graphene flakes on a 300 nm SiO2/Si substrate, illuminated with a white LED, and imaged using a 40×, 0.65NA objective. Graphene flakes, produced via mechanical exfoliation, are widely used to study novel electrical and mechanical properties of 2D materials [14]. Thin graphene flakes, randomly-distributed over the substrate, are commonly identified by manual optical microscopy. Subtle variations in the colour and intensity of transmitted light (≈ 2.3% absorption per layer [15, 14]) can be used to estimate their thickness. This method of identifying flakes is both time consuming and subjective when performed manually, but can be automated with the OFM. Raman spectroscopy measurements have confirmed the flake highlighted in Figure 2b is tri-layer graphene. Unlike other methods of identifying flakes (AFM, Raman spectroscopy, etc.), this searching method is high speed and low-cost, and the graphene is not damaged. By optimising illumination for detection of graphene flakes [16, 17], and improving image processing [14], we estimate the OFM can resolve even monolayer graphene, and we have also imaged the transition metal dichalcogenide molybdenum disulphide with similar success.

#### 2.1.3 Polarisation-contrast imaging

Many microscopic structures important to biology and biochemistry benefit from polarisation-contrast imaging [18, 19, 20]. Anisotropic or chiral structures can rotate the polarisation vector of light propagating through them. The OFM can be used for polarisation-contrast imaging by placing a one linear polariser between the illumination and the sample, and an orthogonal polariser between the tube lens and the sensor (schematized in Figure 2c). Light that passes through the sample unchanged is extinguished, thus only chiral or anisotropic regions of the sample are imaged. Figure 2c shows an image of 4-Cyano-4′-pentylbiphenyl (5CB) liquid crystal droplets, obtained in polarisation-contrast mode with a 40× objective. The sample was prepared by adding a drop of 5CB liquid crystal to the slide, followed by acetone to form a film. The droplets can be seen clearly in the bright-field transmission illumination image, and the liquid crystal orientation structure can be seen in the cross-polarised image. The isotropic background is extinguished, appearing dark. Within the droplets, dark regions appear when the 5CB is aligned parallel or perpendicular to the illumination polarisation, and bright regions appear as the crystal orientation deviates from this.

#### 2.1.4 Fluorescence imaging

Fluorescence microscopy is one of the most important and powerful methods for investigating a range of biological processes. By illuminating a fluorescent sample with suitable excitation-wavelength light, emitted light with a Stokes shift in wavelength can be observed. The OFM can perform low-cost fluorescence microscopy by inserting a dichroic beam splitter and optical filters within a printed filter cube between the tube lens and the sensor, and illuminating with an LED of the desired excitation wavelength (schematized in Figure 2d).

Figure 2d shows fluorescence images of unstained Lily of the Valley (*convallaria majalis*) rhizome taken on the OFM. Two excitation wavelengths (480 nm and 525 nm, with corresponding fluorescence filters as shown in the figure) were used. When illuminating at 480 nm, fluorescence at > 510 nm (green) is imaged, and under 525 nm illumination, fluorescence at > 590 nm (red) is imaged. In principle, by selecting appropriate filters and LEDs, illumination modules can be constructed for any fluorescence wavelength as long as the emission is within the wavelength sensitivity range of the sensor.

### 2.2 Assessing optical resolution and distortion

We characterise the resolution of the OFM by calculating the point spread function (PSF) from a black-white edge in an image. This method does not require nanofabrication techniques or fluorescence imaging capabilities, and is thus appropriate for resource-limited settings. By aligning and averaging several rows of an image with a vertical edge, we recover an edge response function with low noise. Rows are averaged together using a smoothing spline, which provides both noise rejection and sub-pixel alignment. Differentiating this edge response function (after taking the square root to convert from intensity to amplitude) yields a point spread function, with a full width at half maximum of 480 nm for a 40×, 0.65NA Plan-corrected objective. This is very close to diffraction limited performance, over the central 50% of the field of view. Towards the edges of the field of view both field curvature and aberration are visible, resulting in decreased resolution. While this can be alleviated to some extent by acquiring multiple images at different *z* positions, our preferred solution is to tile images together, such that only the central region of each tile is used in the final image, and the lower-quality edges are only required for alignment.

By scanning a sample with straight edge across the field of view it is possible to measure distortion in the optical system. As the optics are axisymmetric, we would expect any distortion also to be axisymmetric, and thus a straight line would appear to deform as it moved across the field of view (appearing straight only in the centre). Performing this experiment with a variety of optical configurations (including an RMS, Plan-corrected objective and a spatially offset Raspberry Pi camera lens), suggested that distortion is very low in this system, less than 0.5%. It is possible that even this slight distortion was affected by the non-uniform response of the Raspberry Pi camera module, and so we suggest it is considered an upper bound.

Calibrating the intensity response of the Raspberry Pi camera module is an important consideration in making the OFM usable. The sensor is designed to be used with a short focal length lens, and consequently suffers from reduced efficiency at the edges of the sensor when used with light that is normally incident across the sensor, such as we have in the microscope. By overriding the default “lens shading table” in the Raspberry Pi’s hardware-accelerated image processing pipeline, we can correct this vignetting, at the cost of increased noise at the edges of the image [21].

### 2.3 Automated imaging

#### 2.3.1 Autofocus

Autofocus is crucial for automated microscopy. As the sample is translated, deviations in the sample thickness, slide angle, and stage position can cause the image to defocus. Manual microscopy can rely on user intervention to correct for this, however for automated microscopy this is not the case. The OpenFlexure Microscope’s software includes two image-based autofocus algorithms, capable of automatically bringing a thin, flat sample into sharp focus.

The first makes use of a Laplacian filter, commonly used for edge detection [22, 23], as a sharpness metric. Additional details of this method are given in Supporting Information 2. The stage is moved sequentially through a set of positions, and at each position an image is captured as a greyscale array, and the sharpness is calculated from this array. While this method is accurate for the vast majority of samples, it is also slow to run on the Raspberry Pi.

The second option is to measure sharpness while moving the stage continuously, by monitoring the size of each frame in the MJPEG video stream. Details of this method are given in Supporting Information 3. This method completes in 3–4 seconds, an order of magnitude faster than the Laplacian filter. The speed and reliability of these autofocus options allow for large tiled images to be captured on the microscope.

#### 2.3.2 Tile scans

One of the many advantages of digital microscopy over traditional microscopy is the ability to automatically obtain large quantities of data over both space and time. These are clearly demonstrated by tiled scanning and time-lapse imaging. Tiled scanning allows images much larger than the normal field of view to be obtained by automatically moving around the sample, taking images at a series of locations, and then digitally reconstructing a complete image. Like panorama photography, some overlap is required between the images, but in principle arbitrarily large scans may be taken within the stage’s range of travel. Unlike traditional panorama photography however, the extremely short depth-of-field associated with optical microscopy presents challenges with staying in focus across the scan. In order to properly recombine tiled images, every image must be properly focused. Thus, the microscope should be brought back into focus before taking each image. Autofocus accounts for a significant fraction of the time taken to make a tiled scan, so the JPEG-based fast autofocus method improves acquisition time by a factor of 5–10.

Figure 3 shows a stitched tile scan of a blood smear sample, obtained from a 10 × 10 grid of captures. The composite image was obtained using Image Composite Editor from Microsoft Research [24]. The figure highlights an individual capture, showing the field-of-view of a single image in the scan. At each *x*-*y* position, the autofocus routine is performed, before taking a *z*-stack of 5 images centred on the focus. The central image of each *z*-stack is used in the stitched image, while the surrounding *z*-positions are used for different analyses of the sample.

**Figure 3:**
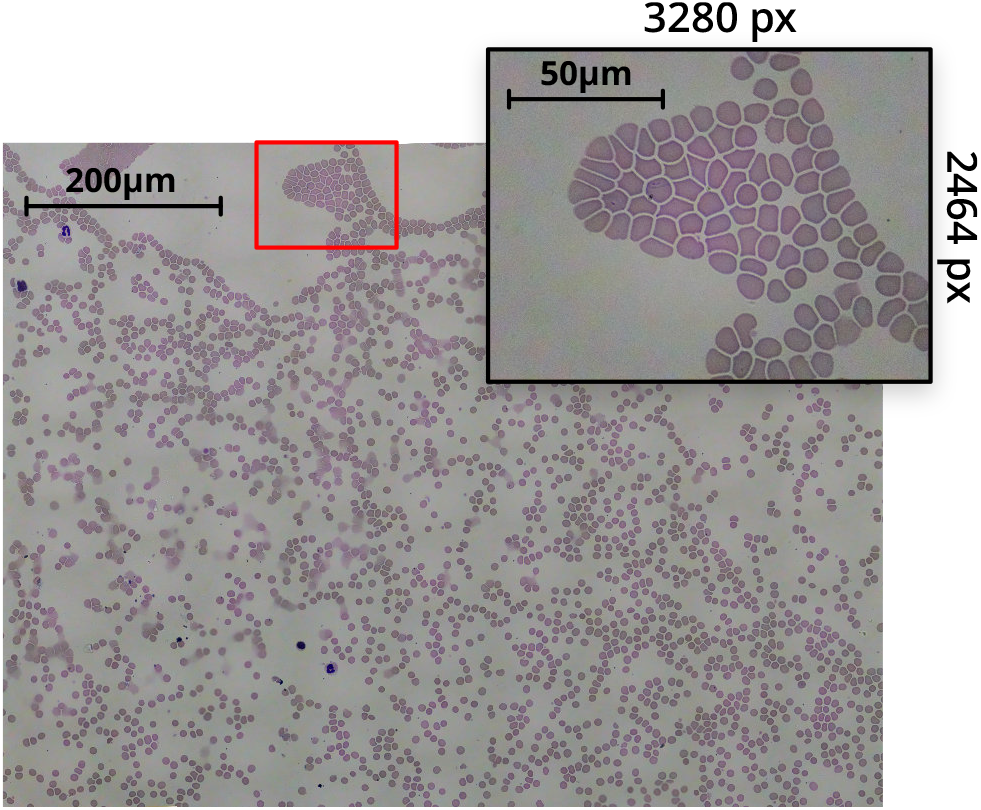
Tiled scan image of a Giemsa-stained thin blood smear, obtained with a 100×, 1.25NA oil immersion objective. The inset highlights an individual 8-megapixel image from the scan. The composite image was obtained from a 10 × 10 grid of captures. After accounting for image overlap and skewing, and cropping out edges of the composite with missing sections, the resulting image is 14920 px ×11270 px (≈ 170 megapixel).

A typical 10 × 10 grid scan with a 5-image *z*-stack at each position (500, 8 megapixel images), running a fast autofocus between each *x*-*y* position, can be obtained in under 25 minutes. Storing raw bayer data per-image increases the acquisition time. By capturing at lower resolutions, a 10 × 10 × 5 scan can complete in around 15 minutes. We have found that depending on where the data is being stored, fluctuations in write-speeds significantly affect acquisition time. However, in future this could be optimised by making better use of parallelisation, reducing the variability in scan acquisition time.

This automation can be used to image samples over long periods of time using time-lapse imaging. Here, the stage position remains fixed, but images are periodically captured. Again, autofocus can be performed before each capture to negate any long-term positional drift. This allows long-term behaviour of samples to be captured in a data-efficient manner, for example, the growth of biological systems over many days or weeks. Slow time-evolution means images need only be captured every few minutes or hours, an impractical and tedious task for a human to perform.

## 3 Software and Usability

Usability and user-experience (UX) have been carefully considered to maximise the breadth of functionality available on the OpenFlexure Microscope. While the microscope software stack is designed to be extensible, aided by full documentation and a comprehensive plugin system, non-developers must be considered the primary users. For this reason, the software is split into a server, running on the microscope itself, and client applications which communicate with the microscope over a network. The server software is distributed as a pre-built SD card image for a Raspberry Pi microcomputer [25], and is common to all OpenFlexure Microscopes.

To ensure stability across a wide range of applications, functionality beyond the basics (stage movement, image captures, and basic hardware configuration) is provided by plugins. Developers are able to create new plugins specific to their needs, and enable them on a per-microscope basis. Crucially, plugins can be disabled or entirely removed from the microscope, allowing for a stable environment to be restored whenever required.

Our primary client application, OpenFlexure eV, is a cross-platform graphical application that both exposes basic functionality, and renders user-interfaces for more complex microscope plugins (tile scanning, autofocus, lens-shading calibration, and more) [26]. The application is designed with a strong focus on general usability and UX, especially for non-programmers. Users of the microscope are able to run comprehensive microscopy experiments, with full control over the format and type of data stored, without any programming experience.

## 4 Distributed Manufacturing and Sustainability

A key aim of the OpenFlexure project is to enable local production. We have demonstrated the local production of microscopes for educational, scientific, and clinical applications at STICLab’s facility in Dar es Salaam, and with our partners at Tech for Trade (Nairobi, Kenya). This has required many optimisations to make the OFM easy to print, assemble, and source parts for, which is equally useful in established research labs. The open-source [27] designs have been replicated in maker spaces and academic labs in numerous countries including in Peru, Germany, Ghana, the USA, and the UK. This replication provides reassurance that the complex printed components can be produced reliably on a wide range of desktop filament deposition printers. This ease of production allows customisation and maintenance of the equipment without external service engineers, and makes the OFM a useful prototyping tool in the development of novel microscopes, or a component to embed within larger instruments.

The 3D printed translation mechanism requires no assembly or alignment. This reduces the bill of materials and speeds up assembly of the microscope. Unlike sliding mechanisms that rely on bearings or smooth ground surfaces, a monolithic flexure mechanism is largely unaffected by dusty or humid environments. Layer-by-layer 3D printing of complicated geometries often requires additional supports structures to be printed, which must be removed by complicated post-print processing. The OFM has been engineered to print without support material, for example by carefully building up unsupported structures using bridges between parts that are present on lower layers, and using a small number of thin “ties” that can be easily and quickly removed during assembly. This makes the design both easier and faster to print, and avoids the risk of damaging printed parts while removing support.

Non-printed parts have been carefully considered to balance cost, performance, and ease of sourcing. Components in the mechanism such as nuts, O-rings, and lenses, push-fit into the design using 3D printed tools. These tools greatly simplify and speed up assembly, allowing each actuator to be assembled in under five minutes. The microscope takes 1–2 hours for an experienced builder to assemble, and typically twice that for a first-time build. Push-fit lens holders ensure that lenses are self-centred, and make it possible to have a single tube extending from the objective to the camera sensor. This sealed tube excludes dust and fungus – a feature introduced in response to trials in The Gambia.

Some components have been carefully chosen to improve the microscope’s lifetime. Previously-used rubber bands perished after weeks to months (regardless of use), and low quality steel screws wore down after 24 hours of continuous use. With quality stainless screws, lubrication, and Viton™O-rings, we are able to run microscopes through around 30,000 cycles of stage motion over several months without failure. Some earlier failures were observed in poorly printed microscopes, or microscopes printed using low quality filament. Nevertheless, in these situations another advantage of 3D printed mechanics comes into play: broken parts can be quickly and easily printed locally and replaced with minimal downtime.

Ultimately, the long-term sustainability of a project such as this one depends on the formation of a community, which is now active on the project’s repositories on GitLab.com. As well as questions and bug reports, we have had contributions with fixes and improvements, for example an adapter to connect the OFM to the “UC2” modular optics system [28], or a rugged enclosure developed by STICLab to protect the microscope in transport and use.

## 5 Conclusions

We have demonstrated that the OpenFlexure Microscope design provides a viable platform for research-laboratory microscopy imaging, owing to both compatibility with high-quality optical components, and robust translation mechanics. The OpenFlexure Microscope design provides precise mechanical sample manipulation in a lightweight and compact device, with comparatively trivial assembly. A range of interchangeable imaging configurations and contrast modes have been developed, suitable for a wide range of applications. We have demonstrated bright-field imaging with both epi- and trans-illumination, as well as polarised light microscopy and multi-channel fluorescence imaging using the OpenFlexure Microscope.

Finally, we have demonstrated the high-speed, automated acquisition of thin blood smears for malaria diagnosis. Microscopes produced in Tanzania have been used in the Ifakara Health Institute to obtain large tile scans of patient sample slides. Ring-form trophozoites of *Plasmodium falciparum* are clearly resolved by the microscope, demonstrating that the quality of images are suitable for diagnosis of malaria in-line with the current “gold-standard”. By combining automated image acquisition with suitable machine-learning, rapid, parallel infection diagnosis is possible, significantly increasing the potential throughput of patients in local clinics.

## Acknowledgements

As the Co-Founder, first CEO, Director of Innovation and Technology Development, and OLI PI at STICLab, Stanley Jaston Mwalembe dedicated his knowledge and scientific skills into contributing to the development of the OpenFlexure Microscope to suit an African context. He keenly saw the real life applications of sayansiScope (OpenFlexure Microscope as branded in Tanzania by STICLab) in schools, colleges, and universities and research centres across Tanzania. His journey and life as a whole have left a remarkable impression on this project, and we will miss his great mind and contributions. May his soul Rest in Eternal Peace. (Stanley Jaston Mwalembe; 11th March 1962-21st March 2019)

We would like to acknowledge financial support from EPSRC (EP/P029426/1, EP/R013969/1, EP/R011443/1), the Royal Commission for the Exhibition of 1851 (Research Fellowship for RWB), and the Royal Society (URF\R1\180153, RGF\EA\181034).

The data that support the findings of this study are openly available in the University of Bath Research Data Archive at https://doi.org/10.15125/BATH-00734.

