## Supporting Information for "Robotic microscopy for everyone: the OpenFlexure Microscope"

### Supplementary Information

#### 1 Estimated cost of production

Costs of common hardware (see Table 1) were obtained from UK retailers, with prices in GBP. For the cost summaries given below, these have been converted to USD at a representative conversion rate of 1.30 USD per GBP.

These estimates do not include time of assembly, which is generally 3-4 hours for an inexperienced builder or 1-2 hours for a more experienced builder, per microscope. Likewise, the cost of 3D printers themselves are excluded.

##### Low-cost version

No motors, cheap webcam as sensor and optics, no microcomputer.

- **PLA:** $\approx$ \$5 USD.
- **Low-cost webcam:** $\approx$ \$5 USD.
- **Common hardware:** Bulk $\approx$ \$5 USD, one-off $\approx$ \$26 USD (see Table 1)

##### Basic version

No motors, Raspberry Pi microcomputer, and Raspberry Pi camera as sensor and optics.

- **PLA:** $\approx$ \$5 USD.
- **Raspberry Pi microcomputer:** \$35 USD.
- **Raspberry Pi Camera rev 2.1:** $\approx$ \$25 USD.
- **Common hardware:** Bulk $\approx$ \$5 USD, one-off $\approx$ \$26 USD (see Table 1)

##### High-resolution, motorised version

Motorised, Raspberry Pi microcomputer, Raspberry Pi camera as sensor, RMS objective and tube lens.

- **PLA:** $\approx$ \$5 USD.
- **Raspberry Pi microcomputer:** \$35 USD.
- **Raspberry Pi Camera rev 2.1:** $\approx$ \$25 USD.
- **Common hardware:** Bulk $\approx$ \$5 USD, one-off $\approx$ \$26 USD (see Table 1)
- **3$\times$ 28BYJ-48 micro stepper motors:** $\approx$ \$5 USD
- **Achromatic tube lens:** $\approx$ \$55 USD.
- **RMS microscope objective:** $\approx$ \$50 USD.
- **Motor driver board:** Varies depending on model. Custom board used in this paper $\approx$ \$50 USD.

| Name | Qty | Supplier | Code | Pack size | Unit price | Pack price |
|---|---|---|---|---|---|---|
| **Common hardware** | | | | | | |
| M3x25mm hexagon head screws | 3 | Westfield Fasteners | WF10920 | 30 | 0.104 GBP | 3.12 GBP |
| M3 brass nut | 4 | Westfield Fasteners | WF20152 | 50 | 0.049 GBP | 2.45 GBP |
| M3 stainless steel washer | 8 | Westfield Fasteners | WF10888 | 200 | 0.009 GBP | 1.8 GBP |
| M3x8mm cap head screw | 14 | Westfield Fasteners | WF2344 | 80 | 0.035 GBP | 2.8 GBP |
| M2x6mm cap head screws | 4 | Westfield Fasteners | WF14259 | 30 | 0.11 GBP | 3.3 GBP |
| $30 \times 2$mm Viton Rubber O-Rings | 3 | Simply Bearings | | 5 | 0.56 GBP | 2.8 GBP |
| **Common electronics** | | | | | | |
| 5mm white LED | 1 | Farnell | 2419079 | 1 | 0.23 GBP | 0.23 GBP |
| 60 Ohm resistor | 1 | Farnell | 2401722 | 10 | 0.0298 GBP | 0.298 GBP |
| **Motorised version** | | | | | | |
| M4x6mm button head screws | 6 | Westfield Fasteners | WF2233 | 40 | 0.06 GBP | 2.4 GBP |

Table 1: Common fastening hardware. Bulk total (using unit prices) 3.80 GBP. One-off total (using pack prices) 19.20 GBP.

#### 2 General autofocus method

The objective is sequentially moved to a range of $z$-axis displacements from the current position. The range, and step size, of this sequence can be freely controlled. At each position, an image is captured as an RGB array, and converted into a greyscale image by taking the mean of the three colour channels at each pixel. A 2D Laplacian filter is applied to the image, highlighting regions of spatially rapid intensity changes (indicative of a well-focused image). The image is squared to prevent regions of positive and negative curvature cancelling out, and the sum of all pixels in the filtered image is then used as a sharpness metric. The objective is returned to the position corresponding to the highest sharpness determined by this metric. While this algorithm generally performs well, capturing RGB arrays and filtering each image in sequence is time consuming.

#### 3 Fast autofocus method

For remote control, the microscope's software serves a real-time MJPEG stream of the camera feed. In this stream, each frame is individually JPEG-compressed. This JPEG compression makes use of the discrete cosine transform. The image is split into blocks, and each block's data can be described as the superposition of 2-dimensional cosine functions of varying amplitude and spatial frequency. Crucially, sharp image features are associated with high-frequency cosine functions. The JPEG compression algorithm will only store amplitudes for the lowest set of frequencies required to accurately represent the image. Therefore, an image containing sharp features across many blocks will require more stored amplitude values, resulting in a larger output file. For this reason, the size of a JPEG image can, with a few key exceptions, be used as a reasonably good metric for overall image sharpness.

For extremely sparse samples, especially when imaged with a contrast mode resulting in a black background, this metric can fail. Far out of focus, more blocks within the image will contain some information. In focus, very few blocks within the image will contain lots of data (high-frequency amplitudes). In extreme cases, the first effect will become dominant and result in a larger overall file size. For a majority of situations studied so far however, this sharpness metric functions well.

The objective first moves to the top of the scan range, then moves down the full scan range in a single continuous movement. Throughout the movement, the position is recorded against time. Concurrently, the file size of each image in the MJPEG stream is stored against time. These two data sets are the correlated to determine an approximate objective position corresponding to maximum JPEG size. The objective is moved back to this position, and the JPEG size measured again. This value is then compared against the curve obtained in the first step to estimate how much further it needs to go, compensating for mechanical backlash. This final movement brings the objective to its target position.
